# Integrated data models on Receptor-Like Kinases for novel domain discovery and functional inference in the plant kingdom

**DOI:** 10.1101/2023.12.21.572927

**Authors:** Qian Liu, Qiong Fu, Yujie Yan, Qian Jiang, Longfei Mao, Long Wang, Feng Yu, Heping Zheng

## Abstract

Receptor-like kinases (RLKs) are the largest signal transduction component in plants, determining how different plants adapt to their ecological environment, resulting in plant-specific ecological niches. Current research on RLKs has focused mainly on a small number of typical RLK members of a few model plants. There is an urgent need to study the composition, distribution, and evolution of RLKs at the holistic level to accelerate the understanding of how RLK assists in the ecological adaptation of different plants. In this study, we have collected 528 plant genomes and established an RLK data model, resulting in the discovery and characterization of 524,948 RLK members. Each member is subject to systematic topology classification and coherent gene ID assignment. Using this data model, we discovered two novel families (Xiao and Xiang) of RLKs. Evolutionary analysis of the RLK families indicates that RLCK-XVII and RLCK-XII-2 exist exclusively in dicots, suggesting that the diversification in RLKs between monocots and dicots could cause differences in downstream cytoplasmic responses. We also use interaction proteome to help empower the data mining of inferring new functions of RLK from a global perspective, with the ultimate goal of understanding how RLKs shape the adaptation of different plants to the environment/ecology. The RLK data model compiled herein, together with the annotations and analytic tools, form an integrated data foundation involving multi-omics data and is publicly accessible via the web portal (http://metaRLK.biocloud.top).

## INTRODUCTION

Adaptation to the environment and ecosystem is essential for the survival and reproduction of biological organisms, requiring the development of a robust signal transduction system to sense external and internal stimuli in animals and plants. While animals primarily rely on G-Protein Coupled Receptors (GPCRs) (Chakraborty and Raghuram, 2023; Hanlon and Andrew, 2015) for signal transduction, the molecular basis of signal transduction in plants follows a different mechanism using Receptor-Like Kinases (RLKs) family of proteins (Shiu and Bleecker, 2001b). The plant kingdom contains more than 300,000 species covering all terrains around the globe despite the lack of mobility, which is only possible with sophisticated and intricate mechanisms to perceive and respond to all environmental conditions. It has become increasingly apparent that RLKs are indispensable in virtually all plant physiological and pathological processes, including cell cycle, growth, development, blooming, and response to environmental stimuli (Shiu and Bleecker, 2001a). Diversity in different ecosystems should also cause considerable evolutionary expansion and diversification of RLKs in different plants, enabling plants to respond precisely to the changing environment of the external world and the coordinated development of various signals between different internal tissue cells (Clark et al., 1997; Clouse et al., 1996; Li and Chory, 1997). While traditional RLK research focuses on limited genes from a few model organisms, studying the composition, distribution, evolution, and function of RLK at the holistic level has become a compelling task.

For decades, it has been thought that most cell-to-cell communication in plants occurs via cytosolic bridges called intercellular filaments. This perception was changed due to the discovery of the first RLK in the 1990s (Walker and Zhang, 1990), which introduced a new paradigm for the mechanism of cell-to-cell communication in plants. Plant RLK is structurally similar to animal Receptor Tyrosine Kinase (RTK), and both contain an extracellular domain (ECD), a hydrophobic transmembrane domain (TM), and a cytoplasmic kinase domain (CD) (Schlessinger, 2014). While the ECD can perceive environmental or internal stimuli produced by organismal cells, the signal can pass along the TM region and trigger the intracellular kinase domain to phosphorylate effector proteins as a common post-translational modification (PTM) phenomenon. Phosphorylation of downstream proteins along the signal transduction pathways can play a variety of functions. For example, it can deactivate or activate proteins, induce protein complex formation, or even guide proper intracellular protein subcellular localization (Day et al., 2016). Due to the high diversity and long-term evolutionary process, RLKs have adapted to various environments. They can participate in a variety of plant growth and immune processes, including stomatal development, meristem development, disease resistance, pollen tube attraction and self-incompatibility, and other life processes. Since the discovery of the first RLK, many RLKs have been characterized from plant cells using molecular biological methods (Walker, 1993).

A holistic understanding of RLKs requires systematic multi-omics analysis, but methodological studies involving RLK lag far behind. The RLK classification scheme currently in wide use by Shiu (Lehti-Shiu and Shiu, 2012) has been proposed for almost two decades and suffers some limitations due to the nature of relying on the information only from the kinase domain. On the other hand, the emergence of second and third-generation sequencing has led the tide of how genetic information related to the RLK is obtained and forms the foundation for the availability of a wealth of RLK-related data. Recent advancement in protein structure prediction, such as Alphafold2 (Jumper et al., 2021), has complemented the known experimental determined RLK structures to make a readily accessible structure repertoire of all RLKs. Nevertheless, most experimental data are not well correlated due to the lack of RLK analytic tools. Even the model plant *Arabidopsis thaliana* has less than 10% of RLK with clearly characterized ligand and interaction networks, let alone other less studied species. While current research on RLK classification and functional annotation mostly focus on a few model plants, the number and diversity of RLKs far surpass what scientists have been able to explore up to date. As more newly-sequenced RLKs remain uncharacterized or unclassified, systematic identification and comprehensive annotation of these proteins become increasingly critical to understand their evolution, biological function, and status in regulatory networks.

Herein, we design and validate a comprehensive RLK data model encompassing more than 500 plant genomes. We also formulate a unified and systematic criterion for the topology classification and gene naming nomenclature for RLKs and RLCKs. Applications of this data model are exemplified by case studies in both novel domain discovery and functional inference: (1) we discover two novel families of RLKs not reported in the literature; (2) we also perform evolutionary and network analysis to implicate the hypothetical function of RLKs subject to further experimental verification. Our data mining and analysis results are publicly accessible through the metaRLK web portal at https://metaRLK.biocloud.top.

## RESULTS

### Identified and unified RLK gene nomenclature

After investigating the current state of research on various plants, we found that some species are less commonly studied and lack a universal naming standard. The gene IDs for less commonly studied species are usually assigned by gene assembly software and lack a correlative meaning with similar RLKs from other species. The lack of a unified naming convention for RLK genes results in chaos for scientists from different research groups to communicate with each other trying to refer to the same RLK. Since an internal ID is mandatory to link data from different sources to carry out all subsequent systematic analysis in a RLK data model, we use this opportunity to craft our internal ID usable as a candidate nomenclature standard. To resolve this dilemma, we formulate and propose the use of a unified gene ID naming norm to standardize the gene ID of RLK in all plants. We use different strategies to assign gene IDs into three classes depending on the availability of commonly accepted gene IDs in the field (Table 1).

**Table 1.** Statistics after renaming the gene ID.

| class | naming convention | genes | species |
| --- | --- | --- | --- |
| I | Preserve original gene IDs in use | 70024 | 55 |
| II | Use gene name from Uniprot | 187 | 9 |
| III | Apply unified gene IDs | 454737 | 473 |

Class I involves species that have been widely studied, such as *Arabidopsis thaliana* and *Oryza sativa*. We keep their gene IDs for 55 species that fit this standard. Class II involves genes not defined in class I but possesses a record in Uniprot (UniProt-Consortium, 2019) that uses the gene ID as the gene name. We take the gene names as the gene IDs of the sequences completely aligned to the records for 187 genes that fit this standard. Class III involves species that are not widely studied and genes that have not been consistently named, encompassing more than 85% of all identified RLKs (Table 1). For such genes, we apply a unified strategy to assign gene ID. We rename these species using a different reference genome. Herein we compare the RLKs of various rice species with *Oryza sativa*, and the RLKs from other species with *Arabidopsis thaliana.* The unified gene name features four parts: (a) abbreviation of the species name; (b) NG/NO/NA part, in which N indicates the chromosome number in which the sequence is located (see details in methods section); (c) the most similar RLK sequence id; and (d) a lowercase letter suffix starting from “a” to distinguish multiple hits most similar to the same gene id in *Oryza sativa* or *Arabidopsis thaliana*.

### Topology classification of RLKs

RLK classification proposed by Shiu (Lehti-Shiu and Shiu, 2012) is based on the homology of the intracellular kinase domain in the cytoplasmic domain (CD) region and disregards the information about the presence and absence of transmembrane (TM) or extracellular domain (ECD) regions. This strategy may be susceptible to insufficient resolution. Another conceptual defect of Shiu’s strategy is that it does not explicitly define the presence of TM region in RLCKs, which could cause the negligence of TM region in many RLCKs. In this study, we formulate a unified and systematic criterion for the topology classification of RLKs considering not only the CD region but also the ECD and TM regions. The boundaries of ECD, TM, and CD regions are defined according to the procedure described in the methods section. Data preprocessing includes a validation step to examine the TM and CD regions. RLK should possess no more than one TM region, with at least a kinase domain in the CD region with a typical length of more than 280 amino acids (Fig. S1A). Therefore, sequences with two or more TM regions or sequences with a CD region shorter than 140 amino acids that indicate incomplete domain structure and compromised kinase activity are excluded from the dataset (Lehti-Shiu and Shiu, 2012) (Fig. S1B), resulting in a validated dataset of 524,948 RLK sequences from 528 plant genomes (Fig. 1A).

**Fig. 1.**
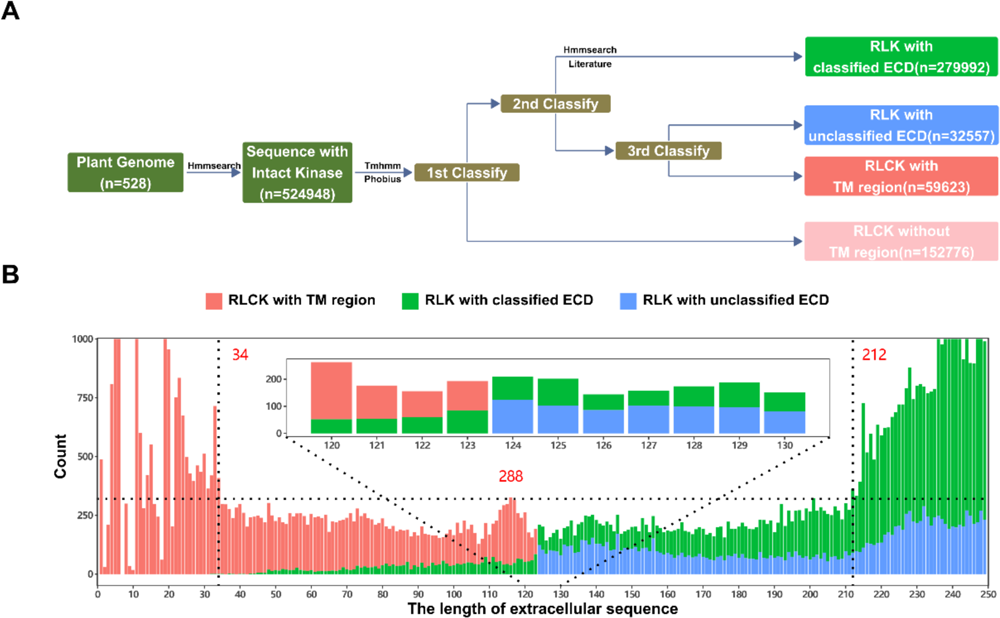
Topological classification of RLK in the dataset. (A) Criterion of topological classification of RLK according to the presence of transmembrane (TM) region and extracellular domain (ECD); (B) Distinguish between RLK and RLCK with TM region based on length of extracellular sequence. RLK with classified ECD colored in green, RLCK with TM region colored in red, RLCK with unclassified ECD colored in blue. The population of RLK or RLCK with TM region are not shown when it exceeds 1000 instances at an ECD length of 5, 6, 11, 19, 239 amino acids.

The validated RLK dataset is subject to systematic topology classification based on three criteria (Fig. 1A). Criterion I identifies sequences with no TM region as ‘RLCK without TM region,’ encompassing 152,776 cytoplasm RLCKs. Criterion II identifies sequences with known ECD as ‘RLK with classified ECD,’ encompassing 279,992 transmembrane RLKs. RLK sequences that fail both criteria I and II are transmembrane proteins without reported ECD, which could either feature a short ECD region and have no ECD (Fig. 1B, colored in red), or possess a potential ECD yet to be characterized (Fig. 1B, colored in blue). Criterion III is used to differentiate between these two cases, with sequences featuring a EC length at or below 123 amino acids representing ‘RLCK with TM region,’ and sequences featuring a ECD length at or above 124 amino acids (Fig. S1C) representing ‘RLK with unclassified ECD,’ encompassing 59,623 RLCKs and 32,557 RLKs, respectively (Fig. 1A). Determination of the classification boundary between 123 and 124 amino acids of extracellular domain is further described in the methods section.

### Structural domain(s) in the ECD region of RLKs

Our RLK topology classification is distinctive to Shiu’s classification, featuring the versatile capability of ECDs to perceive environmental or *in vivo* signals. While the term ECD commonly refers to the full length extracellular sequences in RLK, herein we characterize 780 domains in the ECD region (Table S1), each denote a portion of the ECD region that forms a compact 3D structure, and tag them into 3 subclasses: (a) cECD (Characterized EC Domains) refers to ECDs with determined structure and specific functions that have been studied and described in recent RLK-related literatures (Bender and Zipfel, 2023; Dievart et al., 2020) or the original classification of plant protein kinases by Shiu (Lehti-Shiu and Shiu, 2012) (Table S2); (b) pECD (Pfam Domains not characterized in RLK) refers to many other types of extracellular domains in Pfam (El-Gebali et al., 2019) which have not been reported in RLK-related literatures; and (c) uECD (Unclassified EC Domains) refers to conserved EC sequences of RLKs which forms compact 3D structures with a significant fraction of secondary structure elements but whose functions are yet to be characterized. The validated dataset features the presence of 319,938 cECDs, 74,896 pECDs, and 8,109 uECDs (Fig. 2A).

**Fig. 2.**
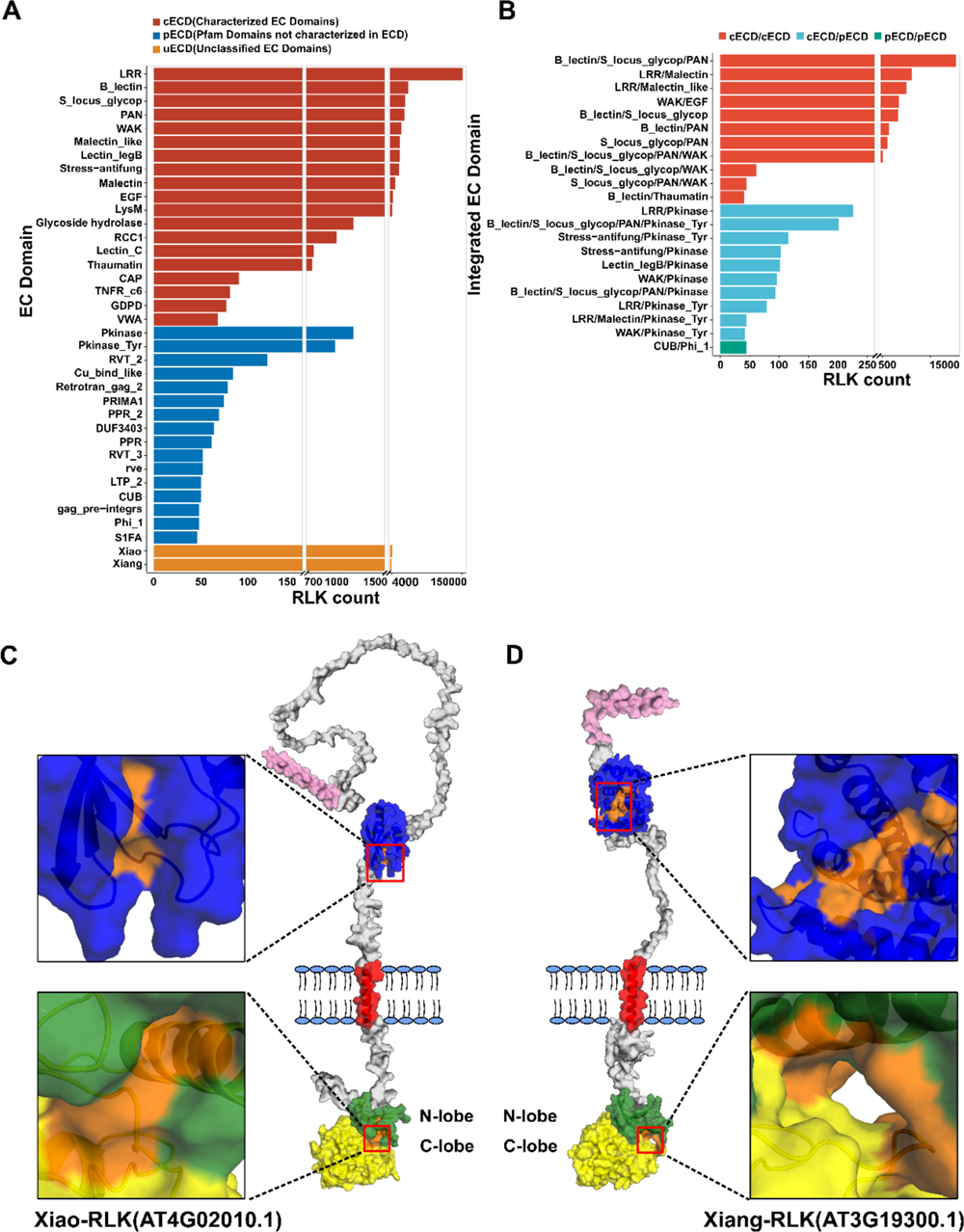
Statistics of RLK extracellular domains from 528 plant species. (A) Statistics on the number of RLK extracellular domains (ECDs) (>40). (B) Statistics on the number of RLK muti-ECDs (>40). (C, D) Two RLK models with new annotated ECDs and their kinase pockets (Pink, blue, red, green and yellow denote signal peptide, the ECD domain, TM region, the N-lobe and the C-lobe of the kinase domain, respectively. Orange denotes the substrate binding sites of the ECD domain or the kinase domain).

Statistics of the most frequently observed cECDs and pECDs indicate that 11 cECDs appear more than 4,000 times in our dataset, while four cECDs and two pECDs appear between 700 and 1,500 times in our dataset. LRR is the most commonly observed cECD, followed by three S-domain cECDs: B-lectin, S-locus-glycop, and PAN. It is worth noting that pECDs include domains that hypothetically sense external stimuli not previously reported to be associated with RLK, such as nucleic acid binding domains (e.g., RVT_2, rve, or S1FA), or metal binding domains (e.g., Cu_bind_like). The addition of either nucleic acid or metal as signal molecules to RLK widens the known repertoire of types of signal molecules that RLK could perceive. Identifying these pECDs opens up an avenue of research that awaits further experimental validation and elucidation. Other specialty pECDs may be used in various other functions, such as PRIMA1 for membrane anchoring, LTP_2 for lipid transfer, or PPR_2 and DUF3403 with unexplored functions (Fig. 2A).

We also systematically analyze all possible multi-ECD combinations and characterize 1,046 combinations (Table S3). The four most commonly observed cECDs, LRR, B-lectin, S-locus-glycop, and PAN, also engage in multi-ECD combinations in the ECD region (Table S2). While LRR is usually partnered with Malectin, Malectin-like, or kinase domain, the other three are S-domain cECDs and make frequent companionships with one another or with the kinase domain (Fig. 2B). The most commonly observed pECD/cECD combination involves pECDs related to phosphorylation (e.g., Pkinase, Pkinase_Tyr) that pair with cECDs (e.g., LRR, S-domain, Stress-antifung) to form dual sequential ECDs (Fig. 2B). Combinative pECDs are less frequently observed, with only one combination CUB/Phi_1 that ranks in the top 20 most populated multi-ECD combinations (Fig. 2B). The functional complexity of multi-ECD combinations implicates that plant may employ these modules to build combinative strategies to sense a series of synergistic, antagonistic, or other spatiotemporal-correlated signals for better environmental adaptation.

### Discovery of novel structural domains Xiao and Xiang in the ECD region

The extracellular sequences of RLK with unclassified ECD from the Alphafold (Jumper et al., 2021) database are clustered using cd-hit (Huang et al., 2010). Clusters with ten or more sequences are further analyzed, resulting in the discovery of two novel ECDs not reported before, namely, Xiao “潇” and Xiang “湘”, symbolizing the ancient Hunan where these two domains are discovered (Fig. 2A). Both newly discovered ECDs exhibit characteristic secondary structural units within the same group. Xiao ECD is similar to the ferredoxin-like domain, which contains a β-strand and two α-helices, such as the *Arabidopsis thaliana* AT4G02010.1 protein (Fig. 2C); while Xiang ECD is similar to the zinc finger domain, it is composed of 6-9 α-helices, such as the *Arabidopsis thaliana* AT3G19300.1 protein (Fig. 2D).

We analyze the differences between the new ECD and the most similar RLK extracellular domains to verify that neither Xiao nor Xiang shares common characteristics with the identified domains. We compare Xiao ECD with several other ECD domains such as Frataxin_Cyay (Gibson et al., 1996) and HMA (Bull and Cox, 1994; Gitschier et al., 1998) domains. We also compare Xiang ECD with several other ECD domains CH (Carugo et al., 1997; Gimona and Mital, 1998; Stradal et al., 1998), OSCP (Wilkens et al., 1997), zf-TAZ (Dames et al., 2002; De Guzman et al., 2000; De Guzman et al., 2004; Goto et al., 2002) and DnaJ_CXXCXGXG (Martinez-Yamout et al., 2000) domains. The result indicates that not only Xiao or Xiang are different in the sequence patterns, but also the two domains don’t contain the heavy metal-binding function, which is a common feature in the aforementioned ECDs. Therefore, Xiao and Xiang represent new extracellular domains of RLK not identified in the Pfam database (El-Gebali et al., 2019). The analysis of newly discovered ECD domains described herein, collectively called uECD (Fig. 2A, Unclassified EC Domains), exemplifies how our RLK data model can be applied to discover novel structural domains.

Fpocket analysis (Peter and Pierre, 2009) shows that both Xiao and Xiang contain small concave pockets that could be connected into larger grooves. We take the largest pockets of Xiao and Xiang ECDs in AT4G02010.1 and AT3G19300.1 of *Arabidopsis thaliana* as an example to illustrate their structural characteristics (Fig. 2C, D). The pocket of Xiao ECD is mainly formed by the coils between α-helix and β-sheet and is composed of two smaller pockets with Druggability Scores of 0.337 and 0.001 as determined by Fpocket. The pocket of Xiang ECD is located at the junction of the α-helices and is composed of four smaller pockets with Druggability Scores of 0.015, 0.001, 0.004, and 0.000. While the scores for individual smaller pockets may not suggest drug binding, their combination forms an extended groove that could potentially bind an elongated polymeric molecule such as peptides.

Xiao ECD contains four subgroups (Table S4). In comparison, Xiang ECD contains seven subgroups (Table S5). The structural features of units between similar subgroups of Xiao or Xiang ECD are evaluated using the Alphafold2 (Jumper et al., 2021) structures corresponding to the domain with the longest sequence in each group as the representative structure. The pairwise structural similarity is calculated using TMscore (Xu and Zhang, 2010; Zhang and Skolnick, 2004) to be in the range of 0 and 1. Xiao ECD exhibits a TM score higher than 0.6 between any two subgroups (Table S4), while Xiang ECD exhibits a TM score higher than 0.15 between any of its two subgroups (Table S5).

Hidden Markov models (HMM) model of these two domains are established for domains in RLKs with unclassified ECD based on all extracellular sequences (Fig. S2, Fig. S3), with the resource downloadable from the metaRLK website. Analysis of Shiu type indicates that Xiao and Xiang are enriched in only a limited number of RLK types. Xiao ECD is enriched in extensin RLKs, 99% of which are “RLK with unclassified ECD,” and about 99% of the extensin sequences contain Xiao ECD. Similarly, Xiang ECD is enriched mainly in RKF3 and URK-1 RLKs. 99% of the RKF3 and URK-1 RLKs are ‘RLKs with unclassified ECD,’ and about 99% of the RKF3 and URK-1 sequences contain Xiang ECD. Though both Xiao and Xiang ECDs are commonly distributed among a wide variety of species, they are not observed in Algae, consistent with Shiu’s description that the three RLK families extensin, RKF3, and URK-1 are not detected in Algae.

### Implication of evolutionary relationship in major branches of Angiosperms

Embarking on a preliminary expedition (Shiu et al., 2004), an evolutionary analysis of the distribution of RLKs has been conducted (Bender and Zipfel, 2023; Dievart et al., 2020) among a lineage of plant species. Herein, we apply the metaRLK data model to investigate the differentiation of two major branches of Angiosperms: monocots and dicots. A typical monocot *Oryza sastiva* and a typical dicot *Arabidopsis thaliana* are selected for comparative evolutionary tree analysis. The results indicate that RLCK-XVII and RLCK-XII-2 families contain only Arabidopsis sequences (Fig. 3A), such as AT5G60080.1 and AT5G60090.1 from the RLCK-XVII family, and AT5G35960.2 and AT5G35960.1 from the RLCK-VI family. Further analysis reveals that these two families exhibit exclusively in dicots (Fig. 3B). The most similar RLCKs found in the corresponding monocots are RLCK-VI and RLCK-XVI, which located at a close monophyletic branch on the evolutionary tree.

**Fig. 3.**
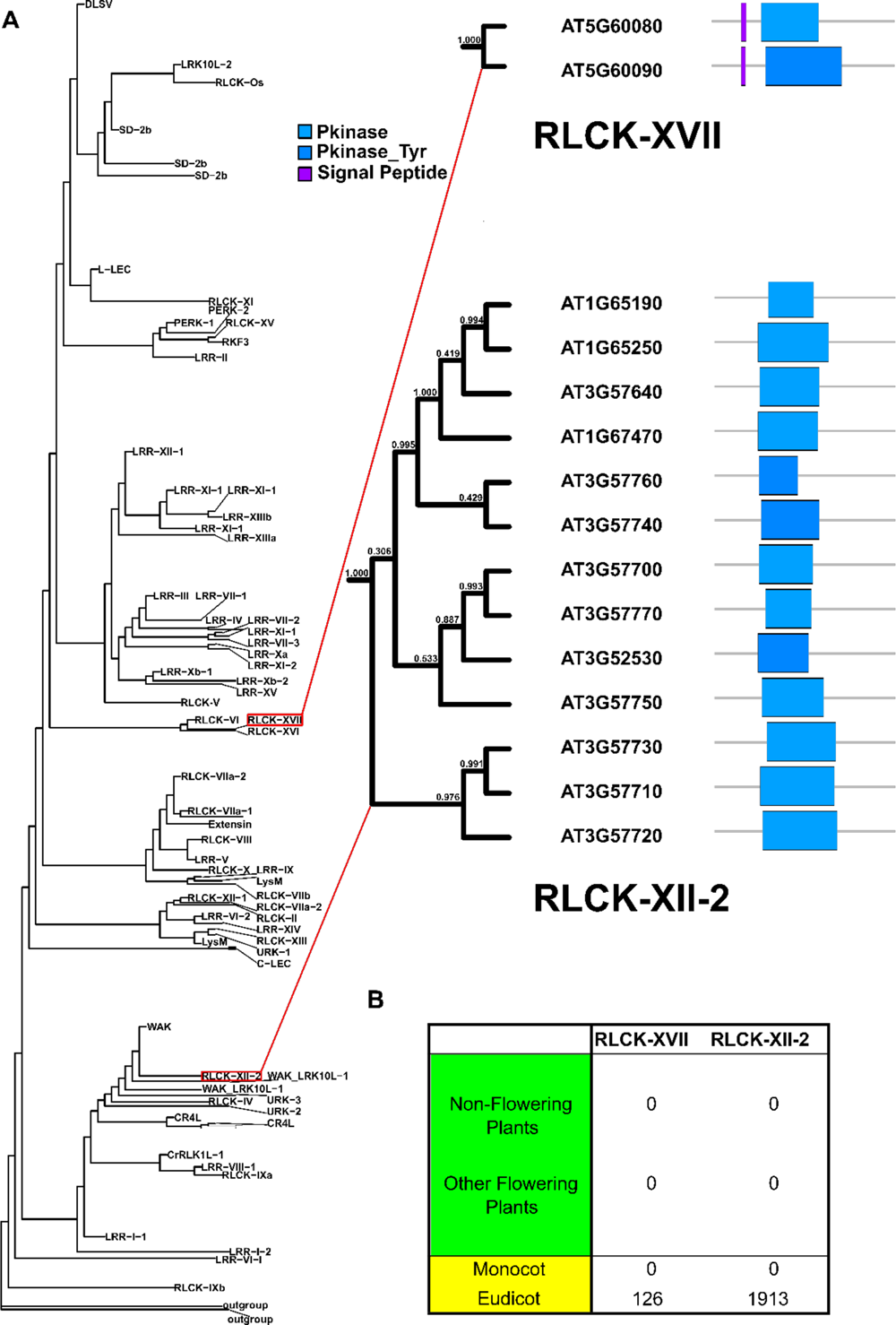
Evolutionary relationship between *A. thaliana* and *O. sativa* RLKs’ kinase. (A) Evolutionary relationship of RLCK-XVII and RLCK-XII-2 families in *A. thaliana* and *O. sativa* RLKs; (B) Distribution of RLCK-XVII and RLCK-XII-2 families in various plants lineages.

Despite of the exclusive presence of RLCK-XVII and RLCK-XII-2 in dicots, the other basic RLK types of rice and *Arabidopsis thaliana* are largely conserved, indicating that the RLKs perform fundamental functions to maintain the vital physiological processes in Angiosperms including both monocots and dicots. The difference in response signals in monocots and dicots could have been achieved through the diversification of the corresponding intracellular pathway along the evolutionary process. Our results illustrate how RLKs in the major branches of Angiosperms (monocots vs. dicots) differentiate in response to signals and adaptation to the environment.

### RLK network analysis for functional inference

Omics-scale analysis of protein-protein interactions (PPIs) plays an important role in understanding the mechanism of RLK signal transduction and furter provide biological insights and testable hypotheses. As we hypothesize that the signaling network resulting from RLK-interacting substrates may be responsible for the functional diversity of monocots and dicots, data about intracellular protein-protein interaction (PPI) are integrated for further analysis. Cytoplasmic substrates interacting with RLK are subject to domain enrichment analysis to discover specific correlation between RLK families and their interacting domains (Fig. 4A). For example, the CR4L family kinase enriches the interacting substrate of Pkinase domain, and the LRR-I-2 family kinase enriches the interacting substrate of C2 domain. Both interacting domains (Pkinase domain and C2 domain) have been reported to interact with the kinase domain of RLK in previous studies (Chen et al., 2023; Du et al., 2016). These results suggest that domain enrichment analysis using cytoplasmic substrates interacting with RLK can identify informative cues to underlying signal transduction mechanisms of certain RLK families, providing clues for quick discovery in RLK functional inference.

**Fig. 4.**
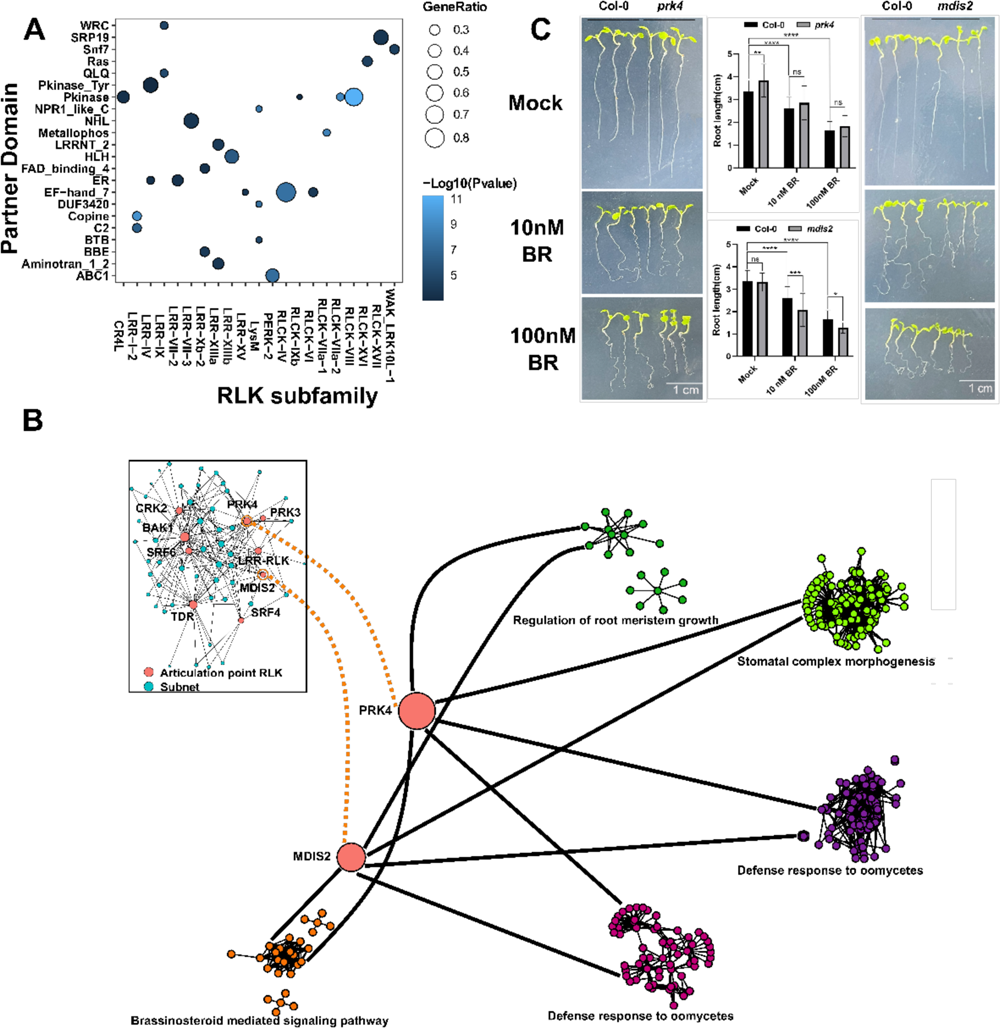
Function inference and intracellular interactions of *A. thaliana* RLK. (A) *A. thaliana* RK intracellular interaction partner domain. (B) PRK4 and MDIS2 articulation point RLK and their subnets which can share new functions in the Arabidopsis thaliana interaction network with them. (C) Phenotypes of *prk4* and *mdis2* mutants under BR treatment.

Species-specific RLK interaction networks are constructed to better use PPI data for RLK functional inference. We screen all known interactions from 177 plant species that have experimentally verified interactions in the String database (Damian et al., 2019). Analysis of the position of RLKs in the interaction network indicates the presence of subnet RLKs that belong to a subnetwork and articulation point RLKs whose removal will result in the disconnection between two or more subnetworks (Fig. 4B) (Smakowska-Luzan et al., 2018). We annotate 1,364 articulation point RLKs and 14,368 subnet RLKs from 87 of the 177 investigated plant species, suggesting that the number of articulation point RLKs is less than that of subnet RLKs. Articulation point RLKs occupy a core position in the protein-protein interaction within a species and regulate the interaction group formed by other proteins, suggesting that such RLKs may play a leading role in modulating mutiple cellular pathway. We hypothesize that the knockout of an articulation point RLK would compromise the coordinating function of the multiple subnetworks to which it connects. To confirm this, we select two articulation point RLKs, Pollen Receptor-Like Kinases 4 (PRK4) (Duckney et al., 2017) and Male Discoverer 2 (MDIS2) (Wang et al., 2016) that perceive female attractant signaling at the pollen, as examples to illustrate the rediscovery of their functions besides double fertilization (Fig. 4B). While the two receptors PRK4 and MDIS2 are verified to enrich in pollen tubes, both of them are also connected to the ‘brassinosteroid (BR) mediated signaling pathway’ subnetwork, suggesting their involvement in the BR signaling pathway (PRK4, P value=0.0000120; MDIS2, P value=0.0000295 (Fig. S4). The hypothetical function is further confirmed experimentally using BR response assay towards knockout mutants of PRK4 and MDIS2. Both PRK4 mutant (*prk4*) and MDIS2 mutant (*mdis2*) (Fig. S4) seedlings grown on BR-containing agar medium exhibit deficiency in growth when compared to the wild-type seedlings (Fig. 4C). While comprehensive understanding of the function implication for a certain RLK requires systematic analysis of all networks, our example illustrates that deciphering the network architecture and understating these topological properties can not only infer the function of uncharacterized RLKs (i.e., RLKs with unknown function), but also rediscover alternative function of characterized RLKs (i.e., RLKs with some reported function).

### Construction of an RLK database and the corresponding web interface

The data and analysis highlighted in this study are compiled to establish a database metaRLK, which fully characterizes a catalog of RLKs from the genomic data of 528 plant species from various resources and formulates the unified and systematic criteria for the topology classification of RLKs, including presence or absence of ECDs, the presence and location of LRR, etc. The data preprocessing workflow includes three modules: (a) Data collection module, (b) Quality control module, and (c) Data processing module (Fig. 5).

**Fig. 5.**
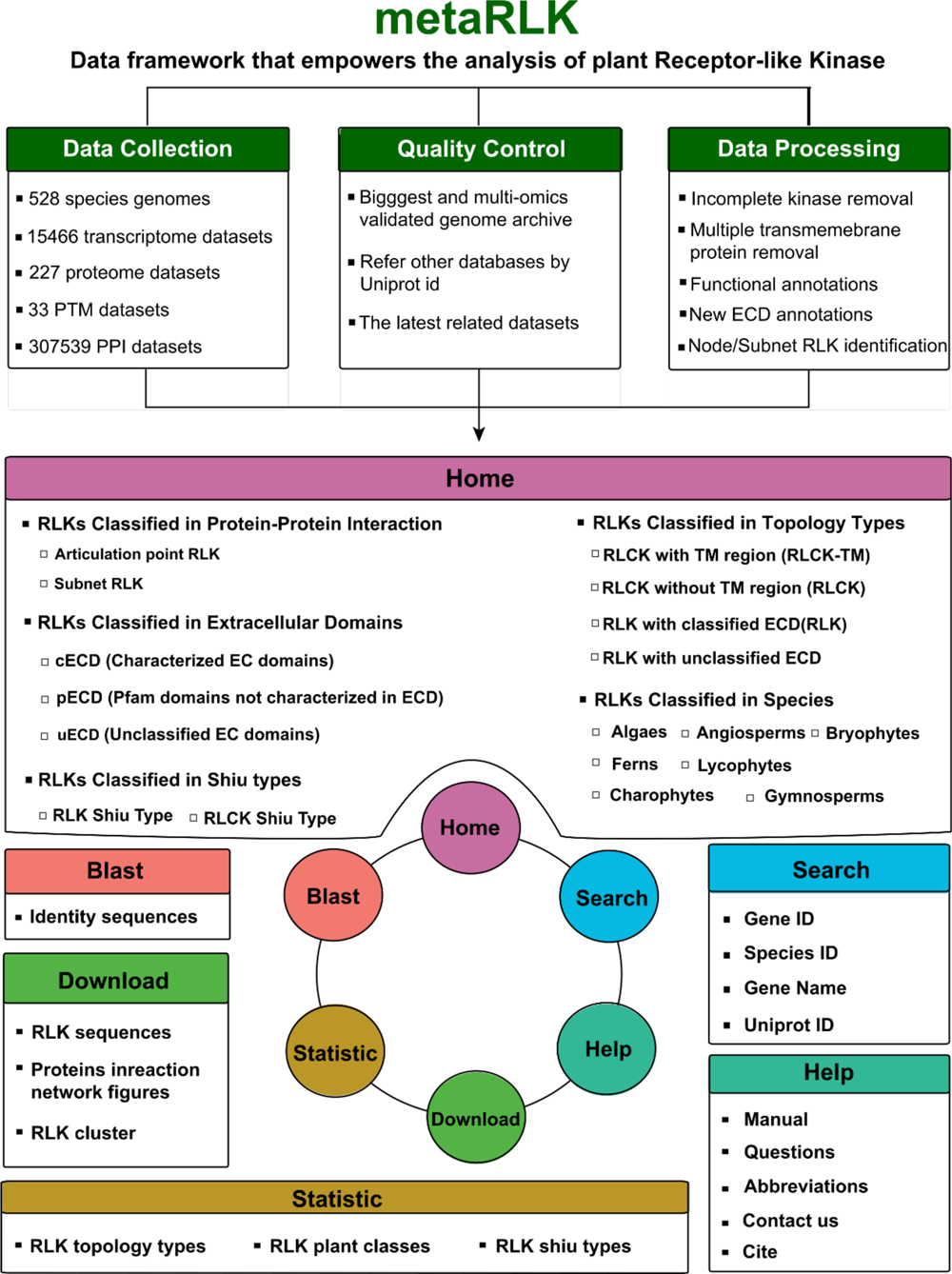
Modules in the metaRLK web portal.

The metaRLK homepage features two summary tables with columns representing the topology classification and rows representing the Shiu classification. The two topology classifications, ‘RLCK without TM region’ and ‘RLCK with TM region’ are mapped to ‘RLCK’ and ‘RLCK-TM’, respectively. The sequences containing uECD are combined with those containing cECD and pECD to form “RLK with ECD” (RLK-ECD). ‘RLK with unclassified ECD’ that do not possess newly characterized ECD are called ‘RLK with intrinsically disordered ECD’ (RLK-IDE). RLK-IDE exists rarely, and their unique extracellular signal transduction mechanism awaits elucidation.

metaRLK provides a framework to annotate a wealth of data from RLK experimental assays, functions, and regulation, including post-translational modification (PTM) data, transcriptomes, proteomes, and protein-protein interaction (PPI) that allows for the network analysis of RLKs to infer its function. The metaRLK is publicly accessible via the web portal located at http://metaRLK.biocloud.top/. This web portal implements six modules: (a) The home module features data summary and entry point for several classification schemes including topology types, Shiu types, species, ECDs, and PPIs; (b) The search module implements several search tools (keywords, sequences, orthogroups, Shiu types) for the quick retrieval of relevant RLK information to aid researchers in locating the RLK of their research interest for further analysis; (c) The Blast module implements sequence tool that use sequence to aid researchers in finding similar RLK of their research interest; (d) The download module allows researchers to fetch RLK data related to specific problem of interest, e.g., The raw data for proteome can be downloaded for further analysis; (e) The statistics module gives an overview of several classification schemes, including the topology types, plant classes, and Shiu types; (f) The Help module includes documentations to help user learn to navigate through the web portal.

## DISCUSSION

Our data mining analysis creates a sophisticated and easy-to-use data model, metaRLK. The verified and well-curated RLK dataset can serve as a benchmark or training dataset to develop a wide range of tools and algorithms for RLK-related analysis to provide biological insights and testable hypotheses. For example, the only dominating RLK classification scheme proposed by Shiu (Lehti-Shiu and Shiu, 2012) is primarily based on the evolutionary homology in the intracellular kinase domain. However, this classification scheme ignores the presence of TM region in RLK sequences, a pivotal feature that determines their subcellular location and implies their function. Many RLCK families defined by Shiu contain a large portion of RLK sequences with TM region (Fu et al., 2022), which is conceptually misleading at first glance. In reality, most of these RLCKs that possess TM region feature a short ECD region and have no ECD domain or uncharacterized ECD domain. In contrast to Shiu’s evolutionary-based approach, we adopt a distinctive strategy to classify RLK and RLCK strictly based on the innate topological differences (Fig. 1A). The topology of an RLK is an essential determinant of its function, and the ability to classify these proteins based on their topology provides critical insights into their roles in signaling pathways.

Comparison between our topology classification and the classification by Shiu (Lehti-Shiu and Shiu, 2012) reveals that the classification scheme proposed by Shiu tends to misclassify sequences that do not conform to the concept of RLK in structure or have only a short extracellular sequence into RLKs. For example, a kinase-containing protein contains multiple TM regions, such as LOC_Os12g26940.1 (Ito and Kurata, 2006; Qiu-min et al., 2004), will be characterized as a RLK by Shiu’s algorithm. Our topology classification treats the raw data prudently using a data preprocessing module to exclude candidate sequences with two or more TM regions. Shiu’s classification also tends to assign proteins with unknown ECD regions as RLK regardless of the length of the ECD region, which causes it to annotate only 48% of the RLCKs characterized by our topology classification procedure. In other words, our topology classification scheme works on a validated RLK dataset that is relatively more prudent and robust to classify RLKs.

Understanding the adaptation of plants to their ecological environment requires a systematic analysis of the RLK-mediated environmental signal response. metaRLK can facilitate the identification of ligands specific to a particular RLK receptor and the characterization of new functions based on RLK network analysis over transcriptomes, proteomes, and interactomes. In addition, our validated RLK dataset enables large-scale determination of extracellular and intracellular protein interaction profiles of RLKs on the genomic and evolutionary levels. Comparative analysis of the similarities and differences in the composition and functional networks of RLKs from more than 500 plant species paves a solid foundation and opens up new avenues toward the data-driven RLK research on plant adaptation to the environment. Our analysis has the potential to facilitate the improvement of agricultural traits based on RLK-mediated signaling pathways in a wide range of applications.

## METHODS

### Source of RLK data from plant genomes

We download genomic protein data of 528 plant species from various resources, including NCBI (O’Leary et al., 2016), Phytozome (Goodstein et al., 2012), GWH (Database Resources of the National Genomics Data Center, China National Center for Bioinformation in 2022, 2022), etc. Then delete the special symbols such as *,?,-at the end and the beginning of the protein sequence, and replace the non-amino acid symbols with X. Finally, use the RLK kinase model (Lehti-Shiu and Shiu, 2012) established by Shiu et al. to identify possible RLK proteins and compare such proteins to the Uniprot database (UniProt-Consortium, 2019). An RLK sequence is mapped to a UniProtKB sequence only if the RLK sequence is 100% identical to the UniProtKB sequence with no insertions or deletions. Otherwise, the closest Uniprot match record will be annotated with the RLK sequence with the corresponding sequence identity.

### Formulation of a nomenclature for the gene ID in RLKs

While the automated structure annotation allows us to classify RLK into different groups from a topological perspective, the function and evolutionary annotation aid in crafting a sensible framework that names homologous RLK of the same group from different species using correlated nomenclatures. A nucleotide sequence is defined as a chromosome or a contig according to the number of proteins with gene names in each plant in the Uniprot database (UniProt-Consortium, 2019) and their ratio to the total number of proteins. The collected chromosome IDs are defined with serial numbers, while all contigs are assigned to the chromosome with serial number 0.

We assign a new gene ID for each RLK in 3 steps: (1) we compare the sequences of each subspecies of rice in metaRLK with the *Oryza sativa* sequence, and after obtaining the most similar sequence, rename the subspecies sequence gene ID in the suffix format of abbreviation +NG+ most similar *Oryza sativa* sequence MSU id suffix, where N represents the pseudo molecule number of the alignment sequence; (2) For RLK sequences not from the rice genus, the same procedure is performed, yet with the comparison object being *Arabidopsis thaliana*; (3) for species that only share the proteome or cannot accurately locate the sequence in the chromosomal position, we change the NG in the naming to NA or NO, where A represents the genome of *Arabidopsis thaliana*, O represents the genome of *Oryza sativa*, and N represents the genome number of *Arabidopsis thaliana* or *Oryza sativa* that is most similar to the aligned sequence. When multiple RLKs from the same species map to the same template RLK, lowercase English letters are added to the end of the gene ID to avoid naming conflict.

### Determination of domains and functional sites

We use the hmmsearch (Finn et al., 2015) program in the HMMER software package to identify domains in the protein sequence according to the domain definition in the Pfam database (El-Gebali et al., 2019), and refer to the annotation results of interproscan (Jones et al., 2014), a non-redundant database developed by EBI that integrates protein families, domains, and functional sites to get annotations of proteins in other databases, as well as *de novo* interpro annotations and GO annotations. Leucine-rich repeat RLK (LRR-RLK, whose ECD region contains LRR domain) is the largest RLK family (Fig. 2A). Plant LRRs have highly conserved characteristic residues and corresponding secondary structures (Appadurai et al., 2019; Kajava, 1998; Kobe and Kajava, 2001), but the boundaries of LRR identified by existing HMM profiles are fuzzy. We adopt a more precise position-specific weight matrix (PSSM) method (Chen, 2021) to locate the boundary of the LRR domains.

### Determination of transmembrane (TM) regions

We use the protocol to characterize TM regions employed by Uniprot(UniProt-Consortium, 2019) with modifications to assign TM regions for RLKs identified from the plant genomes. Tmhmm (Krogh et al., 2001) and Phobius (Käll et al., 2005) were used to predict the TM regions.

Entries predicted with more than three TM regions by both software are excluded from our RLK dataset. For the rest of the dataset, if only a single consensus TM region is characterized, that TM region is used directly. If discrepancies occur between the two software, the prediction result with one TM region takes preference. If more than one TM regions are characterized in both software, such as in the case of LRRC19 (Chai et al., 2009) in which a domain may possess a TM region, the prediction result with fewer TM regions is used, with the TM region immediately prior to the kinase domain being the characterized TM region. A signal peptide region may be assigned if a hydrophobic region is sufficiently close to the N-terminal of the sequence and also possesses other traits of a signal peptide (Krogh et al., 2001). Our TM region assignment procedure results in 69% of the dataset possessing a single TM region, while 31% of the dataset possesses no TM region.

### Determination of the topology classification boundary based on ECD length

The ECD length distribution is further examined to determine the classification boundary (Fig. S1C). A maximal population of 288 RLK sequences is observed at the ECD length of 116 amino acids within the examined region. While populations of the ECD length between 34 amino acids and 212 amino acids do not exceed 288 RLK sequences, many populations of the ECD length outside of this range do. Therefore, the midpoint of these two ends at an ECD length between 123 and 124 amino acids is used as the classification boundary between ‘RLCK with TM region’ and ‘RLK with unclassified ECD’.

### Validation of ECD length

The topology classification scheme is further validated by analyzing the lengths of ECD, TM, and CD for the three classes ‘RLCK with TM region,’ ‘RLK with classified ECD,’ and ‘RLK with unclassified ECD’ (Fig. S5). The lengths of ECD region are largely different among the three classes, with a median of 22 amino acids for ‘RLCK with TM region,’ 405 amino acids for ‘RLK with classified ECD,’ and 252 amino acids for ‘RLK with unclassified ECD.’ Comparison between our topology classification and the classification by Shiu (Lehti-Shiu and Shiu, 2012) using all RLK/Pelle from *Arabidopsis thaliana* indicates that Shiu annotates all RLKs and 335 RLCKs (85%) out of the 409 RLCKs annotated herein. The same analysis performed for RLK/Pelle from all 528 species indicates that Shiu annotates 99.5% of the RLKs and 48% of the RLCKs annotated herein, a good agreement between the two classification schemes.

### Identification of the coding region (CDS) corresponding to the protein sequences

We use seqkit (Shen et al., 2016) to obtain the CDS sequences on the entire genome corresponding to RLK for the 232 species that contain both genomic DNA sequences and protein sequences. Several complications are taken into account to annotate the CDS: (1) alternative splicers could be annotated using transcriptome or proteome data; (2) fusion genes or PTMs could affect gene transcription and translation; and (3) some genomes only annotate the protein corresponding to the longest transcript of the gene, while the other genomes annotate the translation results of all transcripts. We judge the consistency between the translation result of the longest transcript and the protein provided by the researcher and include the CDS sequence that matches the complete transcript. We use genewise of wise2 (Birney et al., 2004) software package to obtain the CDS sequences corresponding to the remaining proteins. The results are combined and presented in the format in which the CDS are separated by introns in the original annotation file.

### Protein structure models

Protein structure models are assigned through the Uniprot (UniProt-Consortium, 2019) match as annotated in Section 1 of the Experimental methods. If the RLK of interest matches a known Uniprot entry, the corresponding protein structure or Alphafold (Jumper *et al*., 2021) model will be retrieved and displayed as the structure model. Static molecular graphics used in the manuscript for demonstration purposes are produced using PyMOL (Schrödinger) (Martinez et al., 2019). Molecular graphics on the view page of the metaRLK web portal use HTML5 as implemented in the NGL Javascript library (Rose Alexander et al., 2019).

### Post-translational modifications (PTM)

Post-translational modifications include phosphorylation sites and palmitoylation sites. For phosphorylation sites, we use protein phosphorylation site annotation for related species in the EPSD database (Lin et al., 2021), and for palmitoyl sites, we use palmitoylation site annotation for related species in the SwissPalm database (Blanc et al., 2019). The positions of PTM sites are visualized using the RCSB-based sequence display plugin (Burley et al., 2023), along with the literature that identifies the site.

### Transcriptomes and proteomes

The mRNA transcription and protein expression levels are integrated for *Arabidopsis thaliana* and *Oryza sativa*. Transcriptome data are retrieved from ExpressionAtlas (Moreno et al., 2022) and PPRD (Yu et al., 2022), while proteome data are retrieved from ProteomeXchange (Deutsch et al., 2023). TPM is used to standardize the expression levels of each sample in the transcriptome, which is convenient for users to compare between different samples. A proteome download channel of the raw data is provided for further analysis.

### Protein-protein Interactions (PPI)

Since the protein interaction databases complement each other, in metaRLK, we integrate the protein interaction information of IID (Chiara et al., 2020), PINA (Cowley et al., 2012), IntAct (Sandra et al., 2014) and String (Damian et al., 2019) databases. The interactions are further annotated using eggnog (Huerta-Cepas et al., 2019), which uses mmseqs pattern to match for a more thorough annotation result to get their functions based on the intersection of GO annotations between the interaction molecules. We also use mcl (Van Dongen, 2008) to subnet division of the protein interaction network (expansion coefficient of 6 to ensure strict division of articulation point proteins) to obtain core RLKs at the ‘Articulation point’ position between two or more subnets, such as PRK4 and MDIS2.

### Construction of RLK Evolutionary trees

After identifying the kinase region of RLK corresponding to RLCK using hmmsearch, we delete the sequence of the domain alignments covering less than 50 percent of the Pfam domain models. For species with alternatively spliced form annotation, only the longest variant of each gene is analyzed further. Putative protein kinases from plants are identified using HMMER v3.0 with Pkinase and Pkinase_Tyr domains. ‘Trusted cutoff’ values specified by Pfam are used as the thresholds.

Evolutionary analysis of the full-length RLK sequences reveals the combination and distribution of different domains to divide RLKs into more representative orthologous groups. RLK sequences are analyzed using Orthofinder (Emms and Kelly, 2019) to evaluate their evolutionary relationship and identify all gene duplication events in those trees. The RLK sequences from each species are used to create a tree to display individual RLK paralogue group by phylo-io.js (Robinson et al., 2016) on the corresponding page from the metaRLK web server.

The protein kinase domain sequences are aligned using muscle (Edgar, 2010), for generating a maximum-likelihood (ML) tree with FastTree (Price et al., 2010) using the CAT model (category approximation of GAMMA model of rate heterogeneity) to account for the rate heterogeneities and the JTT (Jones, Taylor, Thornton) substitution matrix. Evolutionary analysis provides another perspective of tracing the molecular origin.

### Identification of articulation point RLKs and subnet RLKs

Since each interaction database has different methods for judging the credibility of the interaction, here we only divide the experimentally verified part of the String database into articulation point RLKs and subnets. In the process of analyzing the results of mcl (Van Dongen, 2008), we take RLKs that do not belong to any subnets as candidate articulation point RLKs and assume that they may regulate the connected subnets. Furthermore, we selected the RLKs connecting two or more subnets among the candidate proteins as the final results, and each subnet contained three or more proteins.

Gene Ontology (GO) terms validated by the String annotation from the Uniprot database (UniProt-Consortium, 2019) are used to annotate the possible function of articulation point RLKs. To investigate whether some GOs or Pfam domains exhibit articulation point or subnet RLKs enrichment, Fisher’s exact tests are used to determine whether corresponded transcripts ascribed to specific GOs or Pfam domains are more likely to encode RLKs than expected by chance. The calculating formula for the P-value is as follows:

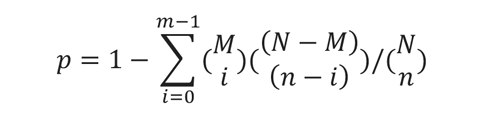

where N is the number of all transcripts with GO or Pfam annotation, n is the number of transcripts containing articulation point or subnet RLKs in N, M is the number of all genes annotated to specific GO or Pfam, and m is the number of transcripts containing articulation point or subnet RLKs in M.

### Functional prediction using articulation point RLKs and subnet RLKs

We systematically generate functional predictions for the articulation point RLKs by identifying functional annotations overrepresented among the interacting subnets related to the articulation point RLKs. These RLKs are classified into different biological process (BP) terms using the BP domain annotations from Gene Ontology (GO). Only the GO terms at the furthest distances from the root term are used to avoid overly generic groupings. Fisher’s exact test (p<0.001) is performed to assess the significance of overrepresented GO categories in an articulation point RLK’s neighbor subnet sets compared to the set of all proteins in the network. Each RLK with an enriched functional annotation in its neighbor subnets is hypothesized to share or have a role in that function.

### Brassinosteroid (BR) response assay

The *prk4* (SALK_020960) and *mdis2* (SALK_112791) T-DNA insertion mutants are obtained from the AraShare Institute (https://www.arashare.cn/index/News/index/cid/15.html). The two mutants are confirmed regarding their T-DNA insertion locations using genomic DNA PCR amplification (Table S6). The exact insertion sites are identified by sequencing (Fig. S4). Seeds are sterilized in 75% (v/v) ethanol grown on half-strength Murashige and Skoog medium (1/2 MS) plates with 0.8% sucrose and 1% phytagel. BR treatment is performed using sterilized seeds on 1/2 MS plates containing 10 nM and 100 nM concentrations of BR. Approximately 30 seeds per genotype are spread onto three individual BR plates per treatment. The seeds are stratified at 4℃ for three days and grown for eight days at 22℃ under long-day conditions (light 16h/ dark 8h). Three biological replicates of were performed with similar results.

### metaRLK database backend and web portal frontend

All data are processed and organized into a PostgreSQL Database Management System. The backend of the metaRLK database uses PostgreSQL 12.11 (database server). The web server is deployed using an Ubuntu Linux virtual machine running Nginx 1.14.0 and Gunicorn 20.0.4. The interface components of the website are designed and implemented using the Django template engine 3.1.4. The position of TM regions and domains are visualized using the RCSB-based sequence display plugin (Burley et al., 2023). metaRLK has been tested in several popular web browsers, including Google Chrome 89.0.4389.82, Mozilla Firefox 87.0, Apple Safari 13.0.2, and Microsoft Edge 89.0.774.75. The styles of the web interface are optimized using the Bootstrap 4.5.0 library to accommodate both large computer screens and small screens on handheld devices. The metaRLK webserver is accessible via https://metaRLK.biocloud.top/.

## AUTHOR CONTRIBUTIONS

H.Z. and F.Y., conceived the project and designed the research.; Q.L., Q.F., Y.Y., Q.J., L.M., and L.W. performed the research, Q.L., Q.F., F.Y., and H.Z. were involved in analyzing the data and preparing the manuscript.

## ACKNOWLEDGEMENTS

The authors would like to thank Dousheng Wu and Yirong Wang for critical reading of the manuscript. The authors would also like to thank Haojie Ma, Yanmin Zhao, Canfen Zhou, and Zhengsan Yu for arranging the virtual machine to set up the web service setup and testing various functionality and compatibility issues of the website.

## REFERENCES

1. Appadurai, R., Uversky, V.N., and Srivastava, A. (2019). The structural and functional diversity of intrinsically disordered regions in transmembrane proteins. The Journal of Membrane Biology 252:273–292.

2. Bender, K.W., and Zipfel, C. (2023). Paradigms of receptor kinase signaling in plants. Biochemical Journal 480:835–854.

3. Birney, E., Clamp, M., and Durbin, R. (2004). GeneWise and genomewise. Genome research 14:988–995.

4. Blanc, M., David, F., and Goot, F. (2019). SwissPalm 2: protein S-palmitoylation database. In Protein Lipidation, (Springer: pp. 203-214.

5. Bull, P.C., and Cox, D.W. (1994). Wilson disease and Menkes disease: new handles on heavy-metal transport. Trends in Genetics 10:246–252.

6. Burley, S.K., Bhikadiya, C., Bi, C., Bittrich, S., Chao, H., Chen, L., Craig, P.A., Crichlow, G.V., Dalenberg, K., and Duarte, J.M. (2023). RCSB Protein Data Bank (RCSB. org): delivery of experimentally-determined PDB structures alongside one million computed structure models of proteins from artificial intelligence/machine learning. Nucleic acids research 51:D488–D508.

7. Carugo, K.D., Bañuelos, S., and Saraste, M. (1997). Crystal structure of a calponin homology domain. Nature structural biology 4:175–179.

8. Chai, L., Dai, L., Che, Y., Xu, J., Liu, G., Zhang, Z., and Yang, R. (2009). LRRC19, a novel member of the leucine-rich repeat protein family, activates NF-κB and induces expression of proinflammatory cytokines. Biochemical and biophysical research communications 388:543–548.

9. Chakraborty, N., and Raghuram, N. (2023). Life, death and resurrection of plant GPCRs. Plant Molecular Biology 111:221–232.

10. Chen, T. (2021). Identification and characterization of the LRR repeats in plant LRR-RLKs. BMC molecular and cell biology 22:1–16.

11. Chen, W., Zhou, H., Xu, F., Yu, M., Coego, A., Rodriguez, L., Lu, Y., Xie, Q., Fu, Q., and Chen, J. (2023). CAR modulates plasma membrane nano-organization and immune signaling downstream of RALF1-FERONIA signaling pathway. New Phytologist 237:2148–2162.

12. Chiara, P., Max, K., and Igor, J. (2020). Informed Use of Protein-Protein Interaction Data: A Focus on the Integrated Interactions Database (IID). Methods in molecular biology (Clifton, N.J.) 2074.

13. Clark, S.E., Williams, R.W., and Meyerowitz, E.M. (1997). The CLAVATA1gene encodes a putative receptor kinase that controls shoot and floral meristem size in Arabidopsis. Cell 89:575–585.

14. Clouse, S.D., Langford, M., and McMorris, T.C. (1996). A brassinosteroid-insensitive mutant in Arabidopsis thaliana exhibits multiple defects in growth and development. Plant physiology 111:671–678.

15. Cowley, Pinese, M., Kassahn, K.S., Waddell, N., Pearson, J.V., Grimmond, S.M., Biankin, A.V., Hautaniemi, S., and Wu, J. (2012). PINA v2. 0: mining interactome modules. Nucleic acids research 40:D862-D865.

16. Dames, S.A., Martinez-Yamout, M., De Guzman, R.N., Dyson, H.J., and Wright, P.E. (2002). Structural basis for Hif-1α/CBP recognition in the cellular hypoxic response. Proceedings of the National Academy of Sciences 99:5271–5276.

17. Damian, S., L, G.A., David, L., Alexander, J., Stefan, W., Jaime, H.-C., Milan, S., T, D.N., H, M.J., Peer, B., et al. (2019). STRING v11: protein-protein association networks with increased coverage, supporting functional discovery in genome-wide experimental datasets. Nucleic acids research 47.

18. Database Resources of the National Genomics Data Center, China National Center for Bioinformation in 2022. (2022). Nucleic Acids Research 50:D27-D38.

19. Day, E.K., Sosale, N.G., and Lazzara, M.J. (2016). Cell signaling regulation by protein phosphorylation: a multivariate, heterogeneous, and context-dependent process. Current opinion in biotechnology 40:185–192.

20. De Guzman, R.N., Martinez-Yamout, M.A., Dyson, H.J., and Wright, P.E. (2004). Interaction of the TAZ1 domain of the CREB-binding protein with the activation domain of CITED2: regulation by competition between intrinsically unstructured ligands for non-identical binding sites. Journal of Biological Chemistry 279:3042–3049.

21a. De Guzman, R.N., Liu, H.Y., Martinez-Yamout, M., Dyson, H.J., and Wright, P.E. (2000). Solution structure of the TAZ2 (CH3) domain of the transcriptional adaptor protein CBP. Journal of molecular biology 303:243-253.

21. Deutsch, E.W., Bandeira, N., Perez-Riverol, Y., Sharma, V., Carver, J.J., Mendoza, L., Kundu, D.J., Wang, S., Bandla, C., and Kamatchinathan, S. (2023). The ProteomeXchange consortium at 10 years: 2023 update. Nucleic acids research 51:D1539–D1548.

22. Dievart, A., Gottin, C., Périn, C., Ranwez, V., and Chantret, N. (2020). Origin and diversity of plant receptor-like kinases. Annual Review of Plant Biology 71:131–156.

23. Du, C., Li, X., Chen, J., Chen, W., Li, B., Li, C., Wang, L., Li, J., Zhao, X., and Lin, J. (2016). Receptor kinase complex transmits RALF peptide signal to inhibit root growth in Arabidopsis. Proceedings of the National Academy of Sciences 113:E8326-E8334.

24. Duckney, P., Deeks, M.J., Dixon, M.R., Kroon, J., Hawkins, T.J., and Hussey, P.J. (2017). Actin–membrane interactions mediated by NETWORKED 2 in Arabidopsis pollen tubes through associations with Pollen Receptor-Like Kinase 4 and 5. New Phytologist 216:1170–1180.

25. Edgar, R.C. (2010). Quality measures for protein alignment benchmarks. Nucleic acids research 38:2145–2153.

26. El-Gebali, S., Mistry, J., Bateman, A., Eddy, S.R., Luciani, A., Potter, S.C., Qureshi, M., Richardson, L.J., Salazar, G.A., and Smart, A. (2019). The Pfam protein families database in 2019. Nucleic acids research 47:D427–D432.

27. Emms, D.M., and Kelly, S. (2019). OrthoFinder: phylogenetic orthology inference for comparative genomics. Genome biology 20:1–14.

28. Finn, R.D., Clements, J., Arndt, W., Miller, B.L., Wheeler, T.J., Schreiber, F., Bateman, A., and Eddy, S.R. (2015). HMMER web server: 2015 update. Nucleic acids research 43:W30–W38.

29. Gibson, T.J., Koonin, E.V., Musco, G., Pastore, A., and Bork, P. (1996). Friedreich’s ataxia protein: phylogenetic evidence for mitochondrial dysfunction. Trends in neurosciences 19:465–468.

30. Gimona, M., and Mital, R. (1998). The single CH domain of calponin is neither sufficient nor necessary for F-actin binding. Journal of cell science 111:1813–1821.

31. Gitschier, J., Moffat, B., Reilly, D., Wood, W.I., and Fairbrother, W.J. (1998). Solution structure of the fourth metal-binding domain from the Menkes copper-transporting ATPase. Nature structural biology 5:47–54.

32. Goodstein, D.M., Shu, S., Howson, R., Neupane, R., Hayes, R.D., Fazo, J., Mitros, T., Dirks, W., Hellsten, U., and Putnam, N. (2012). Phytozome: a comparative platform for green plant genomics. Nucleic acids research 40:D1178–D1186.

33. Goto, N.K., Zor, T., Martinez-Yamout, M., Dyson, H.J., and Wright, P.E. (2002). Cooperativity in transcription factor binding to the coactivator CREB-binding protein (CBP): the mixed lineage leukemia protein (MLL) activation domain binds to an allosteric site on the KIX domain. Journal of Biological Chemistry 277:43168–43174.

34. Hanlon, C.D., and Andrew, D.J. (2015). Outside-in signaling–a brief review of GPCR signaling with a focus on the Drosophila GPCR family. Journal of cell science 128:3533-3542.

35. Huang, Y., Niu, B., Gao, Y., Fu, L., and Li, W. (2010). CD-HIT Suite: a web server for clustering and comparing biological sequences. Bioinformatics 26:680–682.

36. Huerta-Cepas, J., Szklarczyk, D., Heller, D., Hernández-Plaza, A., Forslund, S.K., Cook, H., Mende, D.R., Letunic, I., Rattei, T., and Jensen, L.J. (2019). eggNOG 5.0: a hierarchical, functionally and phylogenetically annotated orthology resource based on 5090 organisms and 2502 viruses. Nucleic acids research 47:D309–D314.

37. Ito, Y., and Kurata, N. (2006). Identification and characterization of cytokinin-signalling gene families in rice. Gene 382:57–65.

38. Jones, P., Binns, D., Chang, H.-Y., Fraser, M., Li, W., McAnulla, C., McWilliam, H., Maslen, J., Mitchell, A., and Nuka, G. (2014). InterProScan 5: genome-scale protein function classification. Bioinformatics 30:1236–1240.

39. Jumper, J., Evans, R., Pritzel, A., Green, T., Figurnov, M., Ronneberger, O., Tunyasuvunakool, K., Bates, R., Žídek, A., and Potapenko, A. (2021). Highly accurate protein structure prediction with AlphaFold. Nature 596:583–589.

40. Kajava, A. (1998). Structural diversity of leucine-rich repeat proteins. Elsevier.

40a. Käll, L., Krogh, A., and Sonnhammer, E.L.L. (2005). An HMM posterior decoder for sequence feature prediction that includes homology information. Bioinformatics 21.

41. Kobe, B., and Kajava, A.V. (2001). The leucine-rich repeat as a protein recognition motif. Current opinion in structural biology 11:725-732.

42. Krogh, A., Larsson, B., Von Heijne, G., and Sonnhammer, E.L. (2001). Predicting transmembrane protein topology with a hidden Markov model: application to complete genomes. Journal of molecular biology 305:567–580.

43. Lehti-Shiu, M.D., and Shiu, S.-H. (2012). Diversity, classification and function of the plant protein kinase superfamily. Philosophical Transactions of the Royal Society B: Biological Sciences 367:2619–2639.

44. Li, J., and Chory, J. (1997). A putative leucine-rich repeat receptor kinase involved in brassinosteroid signal transduction. Cell 90:929–938.

45. Lin, S., Wang, C., Zhou, J., Shi, Y., Ruan, C., Tu, Y., Yao, L., Peng, D., and Xue, Y. (2021). EPSD: a well-annotated data resource of protein phosphorylation sites in eukaryotes. Briefings in Bioinformatics 22:298–307.

46. Martinez-Yamout, M., Legge, G.B., Zhang, O., Wright, P.E., and Dyson, H.J. (2000). Solution structure of the cysteine-rich domain of the Escherichia coli chaperone protein DnaJ. Journal of molecular biology 300:805–818.

47. Martinez, X., Krone, M., Alharbi, N., Rose, A.S., Laramee, R.S., O’Donoghue, S., Baaden, M., and Chavent, M. (2019). Molecular graphics: bridging structural biologists and computer scientists. Structure 27:1617–1623.

48. Moreno, P., Fexova, S., George, N., Manning, J.R., Miao, Z., Mohammed, S., Muñoz-Pomer, A., Fullgrabe, A., Bi, Y., and Bush, N. (2022). Expression Atlas update: gene and protein expression in multiple species. Nucleic Acids Research 50:D129–D140.

49. O’Leary, N.A., Wright, M.W., Brister, J.R., Ciufo, S., Haddad, D., McVeigh, R., Rajput, B., Robbertse, B., Smith-White, B., and Ako-Adjei, D. (2016). Reference sequence (RefSeq) database at NCBI: current status, taxonomic expansion, and functional annotation. Nucleic acids research 44:D733–D745.

50. Peter, S., and Pierre, T. (2009). Fpocket: An open source platform for ligand pocket detection. BMC Bioinformatics.

51. Price, M.N., Dehal, P.S., and Arkin, A.P. (2010). FastTree 2–approximately maximum-likelihood trees for large alignments. PloS one 5:e9490.

52. Qiong Fu, Q.L., Rensen Zhang, Jia Chen, Hengchang Guo, Zhenhua Ming, Feng Yu, Heping Zheng (2022). Large-scale analysis of the N-terminal regulatory elements of the kinase domain in plant receptor-like kinase family. Bioxriv 10.1101/2022.12.10.519927.

53. Qiu-min, H., Hua-wu, J., Xiao-peng, Q., Jie, Y., and Ping, W. (2004). A CHASE domain containing protein kinaseOsCRL4, represents a newAtCRE1-like gene family in rice. Journal of Zhejiang University-SCIENCE A 5:629–633.

54. Robinson, O., Dylus, D., and Dessimoz, C. (2016). Phylo. io: interactive viewing and comparison of large phylogenetic trees on the web. Molecular biology and evolution 33:2163–2166.

55. Rose Alexander, S., Bradley Anthony, R., Valasatava Yana, D.J.M., and Prlić Andreas, R.P.W. (2019). NGL viewer: web-based molecular graphics for large complexes Bioinformatics. Volume 34:3755–3758.

56. Sandra, O., Mais, A., Bruno, A., Lionel, B., Leonardo, B., Fiona, B.-C., H, C.N., Gayatri, C., Carol, C., Noemi, d.-T., et al. (2014). The MIntAct project--IntAct as a common curation platform for 11 molecular interaction databases. Nucleic acids research 42.

57. Schlessinger, J. (2014). Receptor tyrosine kinases: legacy of the first two decades. Cold Spring Harbor perspectives in biology 6:a008912.

58. Shen, W., Le, S., Li, Y., and Hu, F. (2016). SeqKit: a cross-platform and ultrafast toolkit for FASTA/Q file manipulation. PloS one 11:e0163962.

59. Shiu, S.-H., and Bleecker, A.B. (2001a). Plant receptor-like kinase gene family: diversity, function, and signaling. Science’s STKE 2001:re22-re22.

60. Shiu, S.-H., and Bleecker, A.B. (2001b). Receptor-like kinases from Arabidopsis form a monophyletic gene family related to animal receptor kinases. Proceedings of the National Academy of Sciences 98:10763–10768.

61. Shiu, S.-H., Karlowski, W.M., Pan, R., Tzeng, Y.-H., Mayer, K.F., and Li, W.-H. (2004). Comparative analysis of the receptor-like kinase family in Arabidopsis and rice. The plant cell 16:1220–1234.

62. Smakowska-Luzan, E., Mott, G.A., Parys, K., Stegmann, M., Howton, T.C., Layeghifard, M., Neuhold, J., Lehner, A., Kong, J., and Grünwald, K. (2018). An extracellular network of Arabidopsis leucine-rich repeat receptor kinases. Nature 553:342–346.

63. Stradal, T., Kranewitter, W., Winder, S.J., and Gimona, M. (1998). CH domains revisited. FEBS letters 431:134–137.

64. UniProt-Consortium (2019). UniProt: a worldwide hub of protein knowledge. Nucleic acids research 47:D506–D515.

65. Van Dongen, S. (2008). Graph clustering via a discrete uncoupling process. SIAM Journal on Matrix Analysis and Applications 30:121–141.

66. Walker, J.C. (1993). Receptor-like protein kinase genes of Arabidopsis thaliana. The Plant Journal 3:451–456.

67. Walker, J.C., and Zhang, R. (1990). Relationship of a putative receptor protein kinase from maize to the S-locus glycoproteins of Brassica. Nature 345:743–746.

68. Wang, T., Liang, L., Xue, Y., Jia, P.-F., Chen, W., Zhang, M.-X., Wang, Y.-C., Li, H.-J., and Yang, W.-C. (2016). A receptor heteromer mediates the male perception of female attractants in plants. Nature 531:241–244.

69. Wilkens, S., Rodgers, A., Ogilvie, I., and Capaldi, R.A. (1997). Structure and arrangement of the δ subunit in the E. coli ATP synthase (ECF1F0). Biophysical chemistry 68:95–102.

70. Xu, J., and Zhang, Y. (2010). How significant is a protein structure similarity with TM-score= 0.5? Bioinformatics 26:889–895.

71. Yu, Y., Zhang, H., Long, Y., Shu, Y., and Zhai, J. (2022). PPRD: A comprehensive online database for expression analysis of∼ 45,000 plant public RNA-Seq libraries. bioRxiv.

72. Zhang, Y., and Skolnick, J. (2004). Scoring function for automated assessment of protein structure template quality. Proteins: Structure, Function, and Bioinformatics 57:702–710.

